# Harmonic Amplitude-Modulated Singular Value Decomposition for Ultrafast Ultrasound Imaging of Gas Vesicles

**DOI:** 10.1101/2025.07.21.665910

**Authors:** Ge Zhang, Henri Leroy, Nabil Haidour, Nicolas Zucker, Esteban Rivera, Mohamed Nouhoum, Anatole Jimenez, Thomas Deffieux, Dina Malounda, Rohit Nayak, Sophie Pezet, Mikhail G. Shapiro, Mathieu Pernot, Mickael Tanter

**Affiliations:** Physics for Medicine Paris, INSERM U1273, ESPCI Paris, PSL University, CNRS, Paris, France; Iconeus, Paris, France; Institut Curie, Université PSL, Sorbonne Université, CNRS UMR168, Laboratoire Physics of Cells and Cancer, Paris, France; Division of Chemistry and Chemical Engineering, California Institute of Technology, Pasadena, USA; Andrew and Peggy Cherng Department of Medical Engineering, California Institute of Technology, Pasadena, USA; Howard Hughes Medical Institute, California Institute of Technology, Pasadena, USA

**Keywords:** Singular Value Decomposition, Gas Vesicles, Ultrasound Nonlinear Imaging

## Abstract

Ultrafast nonlinear ultrasound imaging of gas vesicles (GV) contrast agents promises high-sensitivity biomolecular visualization with applications such as targeted molecular imaging of tumor markers or real-time tracking of gene expression. However, separating GV-specific signal from tissue remains challenging and requires the implementation of complex transmit schemes. In this work we introduce harmonic amplitude-modulated singular value decomposition (HAM-SVD), which synergizes pulse inversion (PI) with amplitude-modulated singular value decomposition (AM-SVD) to isolate GV-specific second-harmonic signals. In HAM-SVD, single-cycle plane waves at 9.6 MHz and five tilted angles (at a pulse repetition frequency of 2500 Hz) are transmitted under four duty cycles with alternating polarity. Beamformed IQ data are reshaped along a “space × pressure” matrix and decomposed via SVD; tissue background is cancelled by discarding the first and lowest singular modes, yielding an image comprised solely of pressure-dependent second harmonic signals. HAM-SVD sequence enables wide-field, ultrafast imaging without complex transmit sequences. Validation via simulations, *in vitro* phantoms, and *in vivo* rat lower limb experiments demonstrates HAM-SVD’s outperformance compared to PI and AM-SVD. HAM-SVD is shown to achieve a 19.16 ± 1.63 dB signal-to-background ratio (SBR) *in vivo*, surpassing PI (14.19 ± 1.41 dB) and AM-SVD (15.79 ± 1.38 dB). HAM-SVD overcomes limitations of conventional nonlinear techniques (e.g., depth restrictions, tissue clutter) by combining PI’s harmonic sensitivity with AM-SVD’s adaptive clutter filtering of tissue signals. This approach enhances molecular imaging specificity for GVs and holds potential for ultrasound localization microscopy of slow-flowing agents.

## 1. Introduction

Ultrasound imaging plays a pivotal role in biomedical diagnostics, offering non-invasive, real-time visualization with high spatial and temporal resolution [1]. The recent introduction of acoustic reporter genes has further expanded the capabilities of ultrasound, especially in enabling the observation of cellular processes and molecular interactions [2-4]. Among these innovations, gas vesicles (GVs) – air-filled nanostructures encapsulated by a 2-nm-thick protein shell – have emerged as promising biomolecular contrast agents. With typical dimensions of approximately 85 nm in diameter and 500 nm in length [5], GVs exhibit a strongly nonlinear acoustic response under specific pressure conditions (e.g. 0.4 – 0.6 MPa) at a certain transmission frequency (e.g. 15 MHz), generating nonlinear backscattering signals. These nonlinearity can be used to discriminate GVs from tissue signal enhancing detection sensitivity [6]. These properties have spurred the development of nonlinear imaging paradigms, such as amplitude modulation (AM), to exploit GV-specific signals [7, 8].

Conventional nonlinear ultrasound techniques, including parabolic amplitude modulation (pAM) [9] and cross-propagating amplitude modulation (xAM) [7], rely on line-by-line transmissions of three pulses with relative amplitudes of 1/2, 1/2, and 1, in order to achieve GV-specific contrast imaging. However, these methods face inherent limitations in imaging depth and width (due to focus or crossed transmissions), and framerate (due to line-by-line transmission). To address these challenges, Rabut et al. introduced ultrafast amplitude modulation (uAM), which combines AM with multiplane wave transmission and selective coherent compounding [8]. While uAM achieves ultrafast imaging over wider and deeper fields of view compared to pAM and xAM, it requires complex odd- and even-element transmissions to generate 1/2-amplitude pulses. Such odd and even-element transmissions results in the generation of additional deterministic ultrasonic noise that cannot be properly cancelled by nonlinear schemes. This uAM scheme complexity also increases the spatial peak temporal average intensity (I_SPTA_) due to prolonged transmit pulses, even at constant mechanical index. Furthermore, recent work by Nayak et al. integrating harmonic imaging with xAM (HxAM) demonstrated improved GV-specific second harmonic contrast by extracting the second harmonic signals [10]. Nevertheless, the strong need for a simplified transmission scheme combining extended field of view, deeper penetration, fast frame rate and improved SNR still persists.

To streamline ultrafast nonlinear imaging, our group previously developed amplitude-modulated singular value decomposition (AM-SVD) [11], an adaptive contrast-enhanced ultrasound (CEUS) processing technique that integrates singular value decomposition (SVD) with four plane-wave transmissions of varying acoustic pressures (modulated via duty cycle adjustments) at various angles. By exploiting the distinct acoustic responses of tissue, contrast agents, and noise to pressure changes, AM-SVD decomposes acquired frames into singular vectors, and then enabling selective reconstruction of contrast-specific signals with enhanced signal-to-background ratio (SBR). However, AM-SVD operates in the fundamental frequency domain, leaving its potential in second-harmonic imaging unexplored.

Besides, pulse-inversion (PI) imaging is a well-established harmonic technique that transmits pairs of 180°-out-of-phase pulses and sums their echoes to cancel linear components, thereby enhancing second-harmonic generation [12, 13]. Previous literature has shown that, PI could double the harmonic signals, which is not possible with AM [13]. However, conventional PI image scheme has a few limitations. For example, at relatively high mechanical indices, tissue itself also exhibits nonlinear behavior and generate harmonic signals. The overlap of the harmonic signals between tissue and contrast agents can reduce the contrast-to-tissue ratio (CTR), hindering contrast detection. Thus, although PI can effectively capture and enhance the second-harmonic signals, the tissue clutters cannot be removed completely.

To address these gaps, we propose harmonic amplitude-modulated singular value decomposition (HAM-SVD), a novel technique combining the AM-SVD concept with the PI transmission paradigm. HAM-SVD leverages the complementary advantages of both AM-SVD and PI – as PI can effectively capture the second harmonic signals and AM-SVD can effectively filter out the remaining tissue clutter signals that PI transmissions did not completely remove. The objective of this study is to develop and validate HAM-SVD as an ultrafast harmonic imaging technique that improves second-harmonic contrast sensitivity. Through simulations, *in vitro* experiments, and *in vivo* validation, we demonstrate that HAM-SVD achieves superior performance compared to the AM-SVD in suppressing tissue background and enhancing GV-specific second harmonic signals, paving the way for enhanced preclinical and potentially clinical contrast-enhanced ultrasound imaging. Furthermore, HAM-SVD is envisioned to be applicable for the detection of slow moving microbubbles in Ultrasound Localization Microscopy (ULM) [14].

## 2. Materials and Methods

### 2.1. Simulation

Numerical simulations of a tissue phantom with GV contrast were performed in Matlab (MathWorks, USA) to model the harmonic acoustic responses of tissue and GVs, as shown in Figure 1. The relationship between the backscattering amplitude of signals and pressure amplitude (as shown in Figure 1(b)) was extracted from the previous literature [6]. Simulated in-phase quadrature (IQ) data, is basically comprised of three 2D matrices describing the spatial distribution of tissue signal, S_T_, gas vesicle signals, S_GV_, and random noise signals, S_N_, respectively. The backscattering amplitudes of GVs, B_GV_, and tissue signals, B_T_, were simulated with respect to the pressure amplitudes, *p*, according to the relationship demonstrated in Figure 1(a). Further to linear tissue and GV signals, nonlinear tissue signal, B_NL−T_, and second harmonic GV signal, B_NL−GV_, were also simulated based on the previous literature. It was reported that, the second harmonic GV signal is about 5 times smaller compared to its fundamental signal [6]. Therefore, the IQ data in the nonlinear domain can be mathematically expressed as function of spatial and pressure variables:

**Figure 1.**
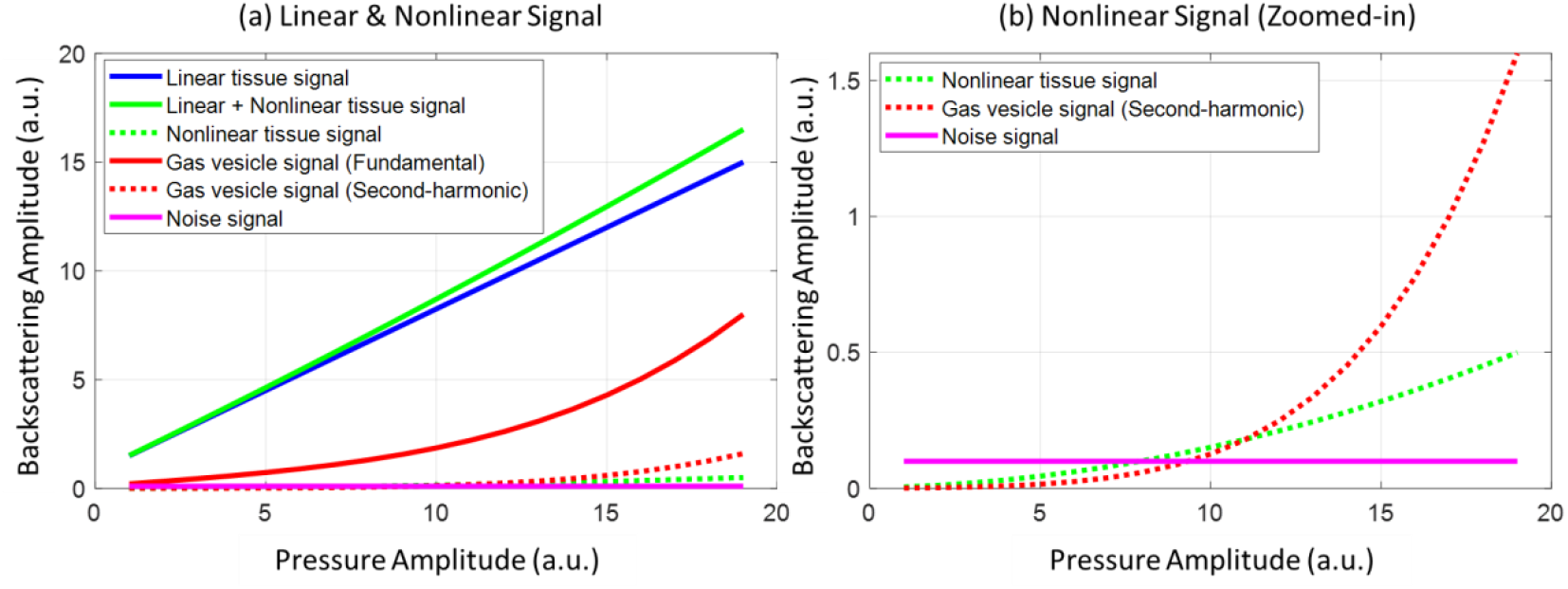
Nonlinear Transmit-receive pressure curves of tissue, GVs and noise for the simulation of backscattering amplitude with respect to acoustic pressure amplitude. (a) Overview of both linear and nonlinear signals from tissue background, gas vesicles, and noise. (b) A zoomed-in comparison between nonlinear tissue signal and second harmonic GV signals.

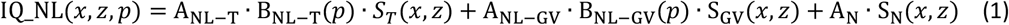

where A_NL−T_, A_NL−GV_, and A_N_ are the scalars which represent the amplitude values of tissue, gas vesicle, and noise components, respectively, in the nonlinear domain. B_NL−T_ and B_NL−GV_ are the amplitude responses to pressure ramp along the curves which demonstrated the relationship between backscattering amplitude and pressure amplitude, *p*, for nonlinear tissue signal and nonlinear GV signals respectively, as shown in Figure 1(b). It has been noted in prior research that tissue may exhibit a nonlinear signal due to the nonlinear propagation of ultrasound waves within tissues, especially at a relatively high acoustic pressure [15]. The nonlinear response of tissue signal was modeled as B_NL−T_(*p*) = *p*^2^, as indicated by the green dotted curve in Figure 1(b), which includes the quadratic term to model the weak nonlinear propagation effects in tissue, as reported in prior studies. The second harmonic GV signal was modeled as B_NL−GV_(*p*) = *p*^5^ + 0.5*p*^4^ + 0.1*p*^2^, as indicated by the red dotted curve in Figure 1(b), emphasizing the higher-order nonlinearity to reflect the pronounced nonlinear scattering of GVs under acoustic pressure, especially in the nonlinear domain.

It can be noticed that, if *A*_*NL*−*T*_ ≫ *A*_*NL*−*GV*_, applying the SVD along the pressure amplitude *p* axis leads to a pressure and space variable separation that can be described as:

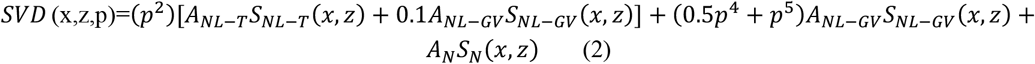

The first term of Eq. (2) corresponds to the first singular vector of the decomposition and contains the weakly nonlinear tissue signals as well as the first-order nonlinearity of GVs. Interestingly, the second singular vector is only related to *S*_*NL*−*GV*_(*x, z*) as can be seen in Equation (3):

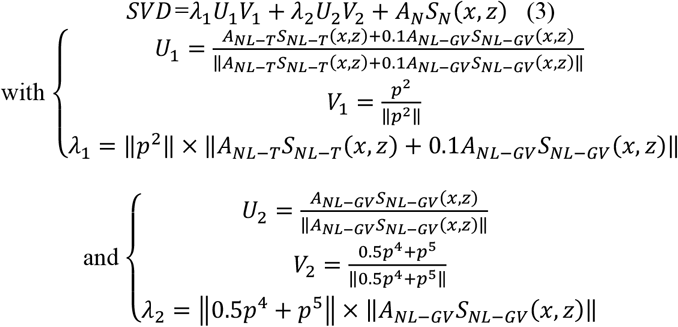

Cancelling the highest singular vector corresponding to weakly nonlinear signals and the lowest singular vectors corresponding to noise will lead to an optimal separation of second harmonic GV signal. The final SVD-filtered image will contain solely the second harmonic signature of GVs and corresponds to:

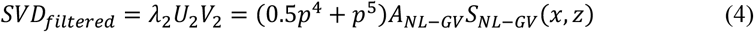

In practice the decomposition of the GV signals can be more complex and spanned over several singular vectors. However, the important point is that SVD decomposition isolated the weakly nonlinear tissue contributions in the first singular vector in such a nonlinear domain. It ensures an optimized extraction of second harmonic GV signals from weakly nonlinear tissue background. Hence, HAM-SVD processing acts as an optimized tissue cancellation filter rather than a GV detection filter. The similar property applies to SVD decomposition along temporal dimension for blood flow extraction [16].

### 2.2. In Vitro Experiment

Nonlinear GVs were prepared following the methodology outlined in Rabut et al. [15]. In brief, GVs were extracted from buoyant Anabaena flos-aquae cells through hypertonic lysis and subsequently purified by repeated centrifugally assisted flotation and resuspension. The outer gas vesicle protein C (GvpC) layer of the GVs was then removed through treatment with 6M urea, followed by an additional round of centrifugally assisted flotation, dialysis in 1x PBS (Slide-A-Lyzer™, 10 kDa MWCO, Thermo Fisher) and resuspension to remove residual GvpC and urea [2].

Single-vial GV phantoms were fabricated by casting 2% (w/v) agarose in PBS supplemented with 0.2% (w/v) Al_2_O_3_ (Sigma-Aldrich). For static imaging, custom 3D-printed molds were employed to create cylindrical wells with a 2 mm diameter. GVs were incubated at 60 °C for 1 minute, mixed in a 1:1 ratio with molten agarose, resulting in a final gas vesicle concentration equivalent to an optical density (OD) of 3.0 at 500 nm (OD_500_ nm), a standard spectrophotometric measurement used to quantify particle concentration in suspension, and loaded into the phantom. The Al_2_O_3_ concentration was carefully selected to match the scattering echogenicity of the gas vesicle well. For the single-vial GV phantom, the GV inclusion was precisely centered at a depth of 9 mm.

### 2.3. In Vivo Experiments

All procedures were performed on a single 7-week-old male Sprague Dawley rat as part of an ongoing project (Project #35411-2022021121597058 v4). The experiment presented in this article was conducted before euthanasia. The experiments complied with French and European regulations and were approved by the local ethics committee (Comité d’éthique en matière d’expérimentation animale, number 59, Paris Centre et Sud). The rat was housed in a group of three per cage under a 12-hour light/dark cycle, at a constant temperature of 22°C, with a humidity level maintained between 45% and 50%. Food and water were provided ad libitum. Before the experiments, the animals were given a minimum acclimatization period of one week to adjust to the housing conditions. All experiments adhered to ARRIVE guidelines and relevant animal care regulations.

Anesthesia was maintained with 2% isoflurane (delivered via a nose mask, Minerve, France). The eyes were protected with moisturizing gel (Ocry-Gel, Virbac, France). Body temperature was maintained at 37°C using a heating bed, monitored by a rectal probe (Physitemp, Clifton, USA). Heart rate and respiratory rate were monitored using a PowerLab data acquisition system with LabChart software (ADInstruments, USA). The imaging was performed using a L15 transducer with the pulse sequence described below in the data acquisition. For imaging of GVs in the lower limb of the rat, the rat was placed in a prone position, with the ultrasound transducer positioned on the left lower limb tissue. Prior to imaging, GVs were mixed with 42 °C 2% agarose solution for a final GV OD_500nm_ equal to 3. An 18-gauge needle was filled with the mixture of agarose and GVs. The mixed solution (0.05 mL) was injected at the depth of 6 mm in the left lower limb of the rat.

### 2.4. Harmonic Amplitude-Modulated Singular Value Decomposition

The detailed imaging acquisition and processing pipeline of HAM-SVD can be seen in Figure 2. It should be noted that, the proposed SVD method utilizes the second harmonic GV signals, as it exploits the pressure dependent nonlinear response of GVs and tissue at the second harmonic frequency. A detailed calculation of theoretical frame rate analysis can be found in Supplementary material 1.

**Figure 2.**
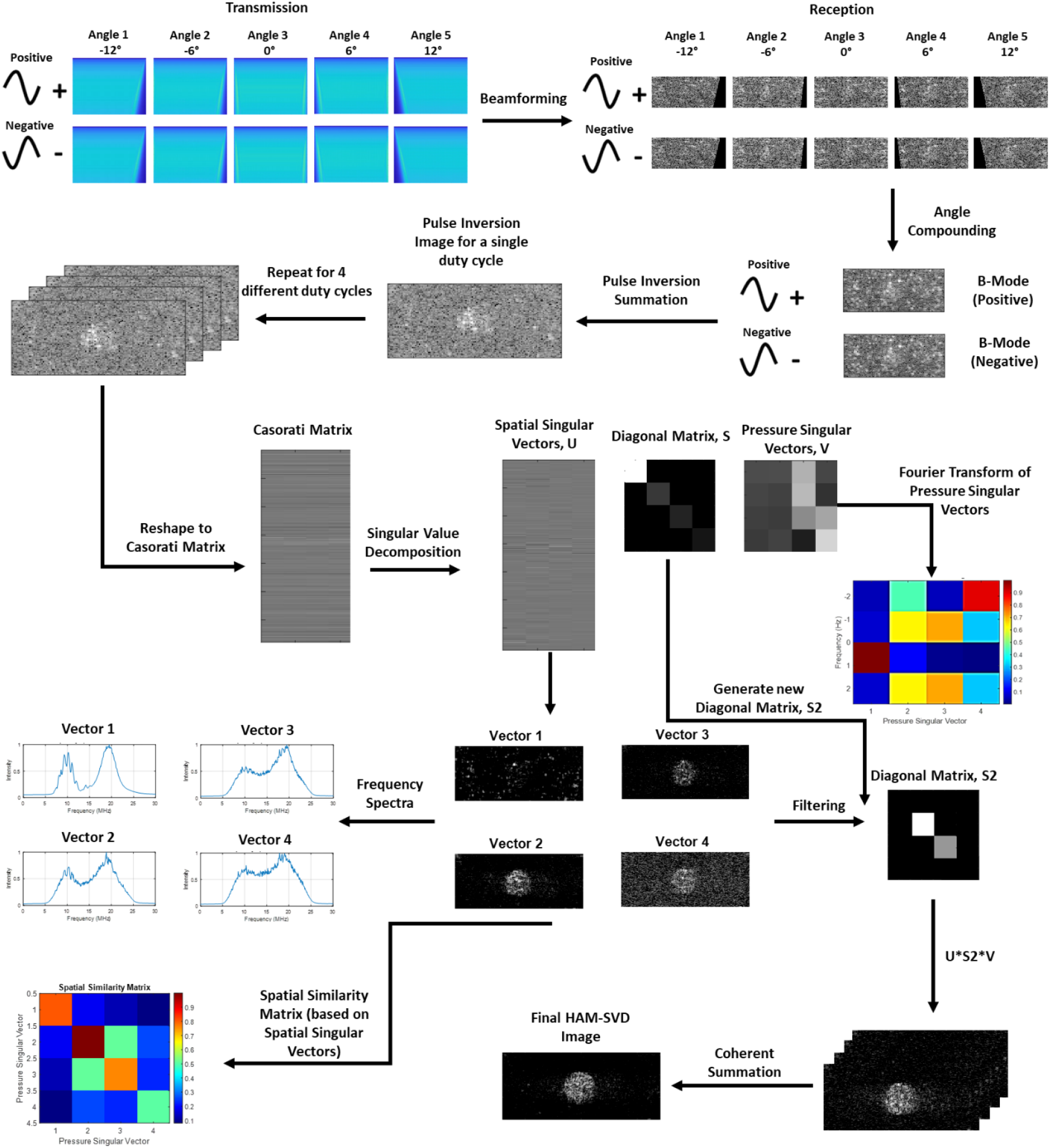
Acquisition and processing pipeline of harmonic amplitude-modulated singular value decomposition (HAM-SVD). For the transmission, positive pulses were transmitted at five different angles and followed by phase-inverted pulses with the same magnitude at the same angles. The received RF data was then beamformed for each angle and then angular coherent compounding was performed to obtain the corresponding positive and negative compounded frames. Pulse inversion summation was performed to generate the second-harmonic image. The above processes were repeated for 4 times with varying duty cycles. Then the resulting four pulse inversion images were reshaped into a Casorati matrix to perform singular value decomposition. After SVD, spatial singular vectors, U, can be displayed to see the performance of decomposition. At the same time, the frequency spectra can be displayed based on the Fourier transform of each spatial singular vector, U. Additionally, spatial similarity matrix can be generated to distinguish the separation among tissue background, contrast signals, and noises. Besides this, Fourier transform of the pressure singular vectors, V, can also helped with the SVD threshold selections. After SVD thresholding, the original diagonal matrix, S, was filtered to generate a new diagonal matrix, S2. The new S2 was used to regenerate the filtered dataset for the corresponding four varying duty cycles. Finally, the images were coherently summed to generate the final HAM-SVD image.

For image acquisition, ultrasound sequences were implemented and executed on a research ultrasound system (Verasonics, USA) driving a linear ultrasound transducer (128 elements, 15.625 MHz central frequency and a 67% bandwidth at -6 dB). All acquisition scripts and processing codes were developed in Matlab (Matworks, USA). For the HAM-SVD pulse sequence, single-cycle plane waves were transmitted at a frequency of 9.6 MHz, with 5 compounded angles for each pulse with an angle range from -12° to +12°. Then the pulses were repeated for 4 different duty cycles and opposite polarities (+1 and -1). The duty cycle (DC) is defined as the ratio of the pulse duration to the pulse repetition period, representing the fraction of time the transducer is actively transmitting during each cycle. Higher duty cycles result in greater energy transfer, which increases the effective average intensity of the transmitted wave. By varying the duty cycle, the transmit amplitude is modulated, enabling the acquisition of nonlinear responses at different pressure levels. The ultrasound raw radio frequency (RF) data was acquired and stored for later image beamforming.

For image beamforming, the RF data were offline beamformed using a delay-and-sum beamformer on GPU, utilizing a resolution grid with a spacing of 0.5λ, resulting in the generation of IQ data for each angle and pulse polarity at the corresponding duty cycles. Then the angle compounding was performed for the positive and negative pulse polarities correspondingly. After that, pulse inversion summation was performed to generate the pulse inversion image at the corresponding duty cycles. As can be seen in Figure 2, four different duty cycles were used in the demonstration.

For image processing, four PI images at different duty cycles were reshaped into a Casorati matrix for singular value decomposition processing. After the decomposition, the spatial singular vectors, U, pressure singular vectors, V, and the diagonal matrix, S, were obtained for SVD threshold selection. Frequency spectra of spatial singular vectors can be used to visualize the distribution of fundamental and second-harmonic frequency components. Spatial similarity matrix based on the spatial singular vectors can be used to observe the spatial coherence between each spatial singular vector. Fourier transform of the pressure singular vectors can be used to visualize the variation in frequency components along the pressure singular vectors. Similar spatial-spectral SVD decomposition was recently applied to brain tissue displacement with an automatic selection of the singular vector corresponding to a physiological rhythms [17]. The detailed procedure of frequency spectrum, spatial similarity matrix, and Fourier transform of the pressure singular vectors were described in the next section. After the threshold selection, a new diagonal matrix, S2, was used along with the matrices, U and V, to regenerate the filtered image dataset at four different duty cycles. Finally, these filtered images were coherently summed in order to generate the final HAM-SVD image.

### 2.5. Evaluation on Singular Vectors

In this study, three methods (Fourier transform of pressure singular vectors, spatial similarity matrix, and frequency spectrum of spatial singular vector) were used to study the spatial singular vectors, U, and the pressure singular vectors, V. These three matrices display more comprehensive and direct visualization of the differences of spatial and pressure singular vectors for each mode after singular value decomposition.

Fourier transform (FFT) of pressure singular vectors was performed to evaluate the change in frequency at each pressure singular vectors. The Fourier transform of pressure singular vectors converts the pressure-amplitude domain signals into a pressure-frequency domain, revealing the spectral characteristics of the underlying components (tissue, contrast, or noise). Each pressure singular vector represents the pressure evolution of a specific mode. Its Fourier transform shows the dominant frequencies associated with tissue motion (low-frequency), contrast (broad band), noise, and also the energy distribution across frequencies for each mode.

Spatial similarity matrix (SSM) was performed by calculating the correlation between all pairs of normalized spatial singular vectors. Tissue and contrast subspaces can be observed from the SSM figure. A high-correlation block (top-left square) normally corresponds to tissue-dominated singular vectors, as tissue exhibits strong spatial coherence. A lower-correlation block represents contrast-dominated singular vectors, since contrast signal introduces spatial-pressure decorrelation, reducing pairwise correlations. The off-diagonal regions demonstrates the weak correlations (darker areas), which reflect dissimilarity between tissue and contrast subspace.

Frequency spectra of spatial singular vectors were obtained by firstly performing SVD on the corresponding RF data at four different duty cycles. Then, FFT was applied on the spatial singular vectors which obtained from SVD to generate the corresponding frequency spectrum for each spatial singular vector.

### 2.6. Image Evaluation

The parameter used for the evaluation of the image quality was signal-to-background ratio (SBR) evaluating the GV contrast as previously described in the literature [18]:

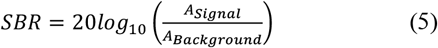

*A*_*Signal*_ is the mean amplitude of signal within the contrast region; *A*_*Background*_ is the mean amplitude of signal within the tissue region.

## 3. Results

### 3.1. Frequency Component Analysis in Original B-Mode and Pulse Inversion Image

Figure 3 illustrated how B-mode and pulse-inversion (PI) imaging capture GV contrast and the corresponding frequency spectra across a range of peak positive pressures (0.067-0.534 MPa), and how applying a second-harmonic bandpass filter affects the imaging. It can be seen from the figure that, after the peak positive pressure of around 0.1 MPa, GV contrast started to be observed in pulse inversion images. After the peak positive pressure of around 0.3 MPa, GV contrast started to gradually decrease. These were also reflected on the frequency spectra, as the peak of second harmonic frequency components significantly increased from 0.1-0.3 MPa. As the second-harmonic frequency component increases, more contrast can be observed on pulse inversion imaging. Although the second harmonic bandpass filter can help B-mode image to significantly improve the GV contrast, however, the noise floor was still higher than that on original PI images without applying the bandpass filter.

**Figure 3.**
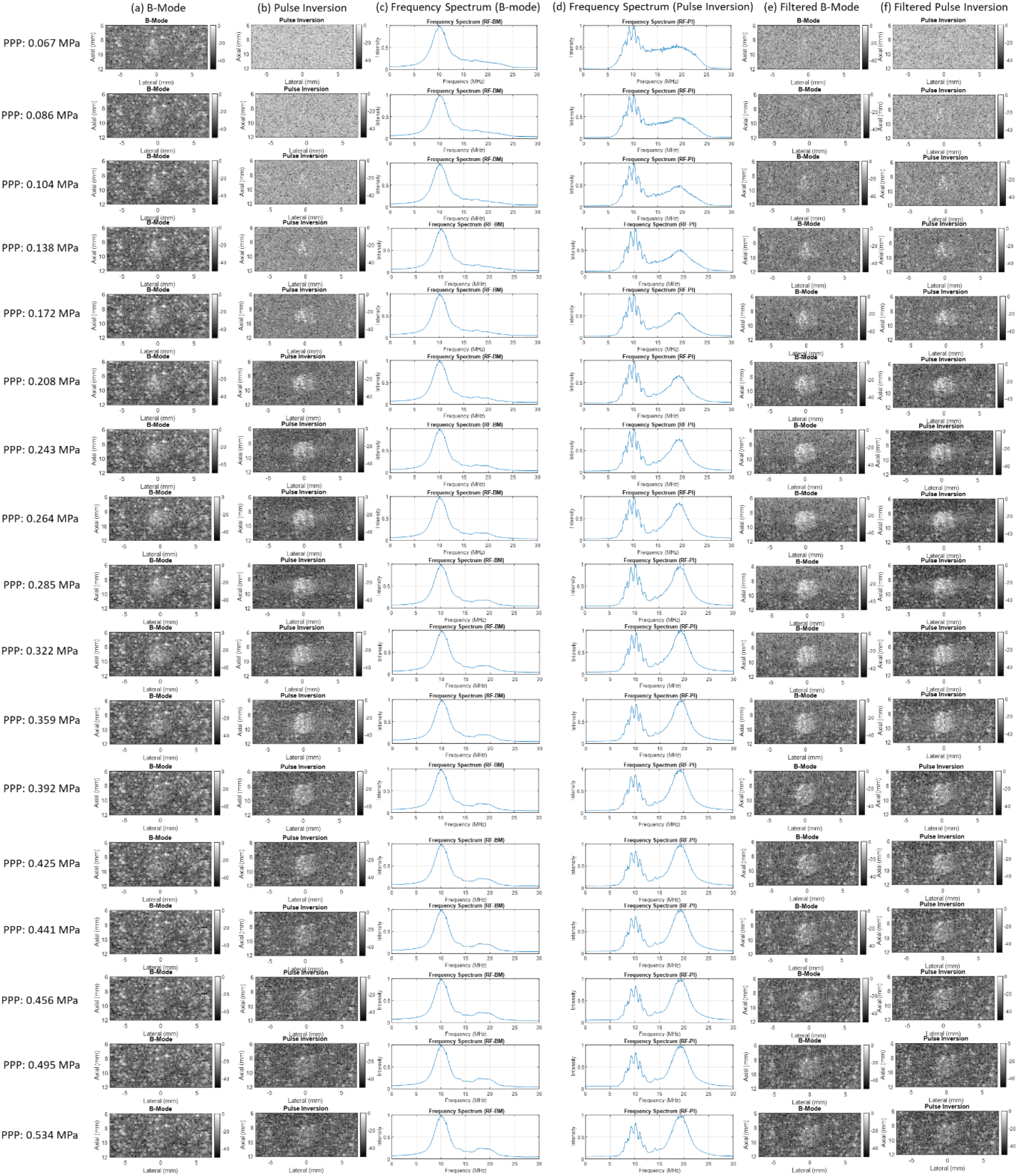
(a) B-mode and (b) pulse inversion images acquired at a transmission frequency of 9.6 MHz and (c-d) their corresponding frequency spectra and (e-f) the corresponding images after the second-harmonic frequency filtering across a peak-positive-pressure range from 0.067 MPa to 0.534 MPa.

### 3.2. SBR Quantification

Figure 4 summarizes how fundamental and second-harmonic frequency components from GVs evolve with increasing acoustic pressure, and how these changes correlate with SBR for conventional B-mode and PI imaging. Figure 6(a) showed the fundamental frequency magnitude rises approximately linearly, as peak positive pressure increases from 0.067 MPa to 0.534 MPa. Figure 6(b) showed the second-harmonic frequency magnitude increased nonlinearly over the same pressure range. To focus on GV response before collapse, Figure 6(c-d) restrict the pressure range to 0.067-0.285 MPa. Figure 6(c) showed the fundamental frequency magnitude increased as the rising pressure before GV collapse for both B-mode and PI imaging. The gradient of PI response was smaller than that of B-mode response. Figure 6(d) showed the response of second-harmonic frequency magnitude before the GV collapse for both B-mode and PI imaging. It can be seen that, PI imaging demonstrated a high degree of nonlinearity of second harmonic frequency magnitude as the pressure increased compared to B-mode imaging. Figure 6(e) showed the ratio between fundamental and second-harmonic frequency magnitude as a function of peak positive pressure for B-mode and pulse inversion images. Figure 6(f-g) showed that SBR quantification for B-mode and pulse inversion images without and with second-harmonic frequency filtering. It can be seen that, the highest SBR that B-mode and PI images can achieved were 5.24 ± 1.46 dB and 12.24 ± 1.65 dB. After the second harmonic frequency filtering, the highest SBR that filtered B-mode and PI images can achieve were 11.27 ± 1.56 dB and 13.10 ± 1.87 dB. Figure 6(h) showed the SBR has a positive correlation with fundamental frequency component on the B-mode image and with second harmonic frequency component on the PI image.

**Figure 4.**
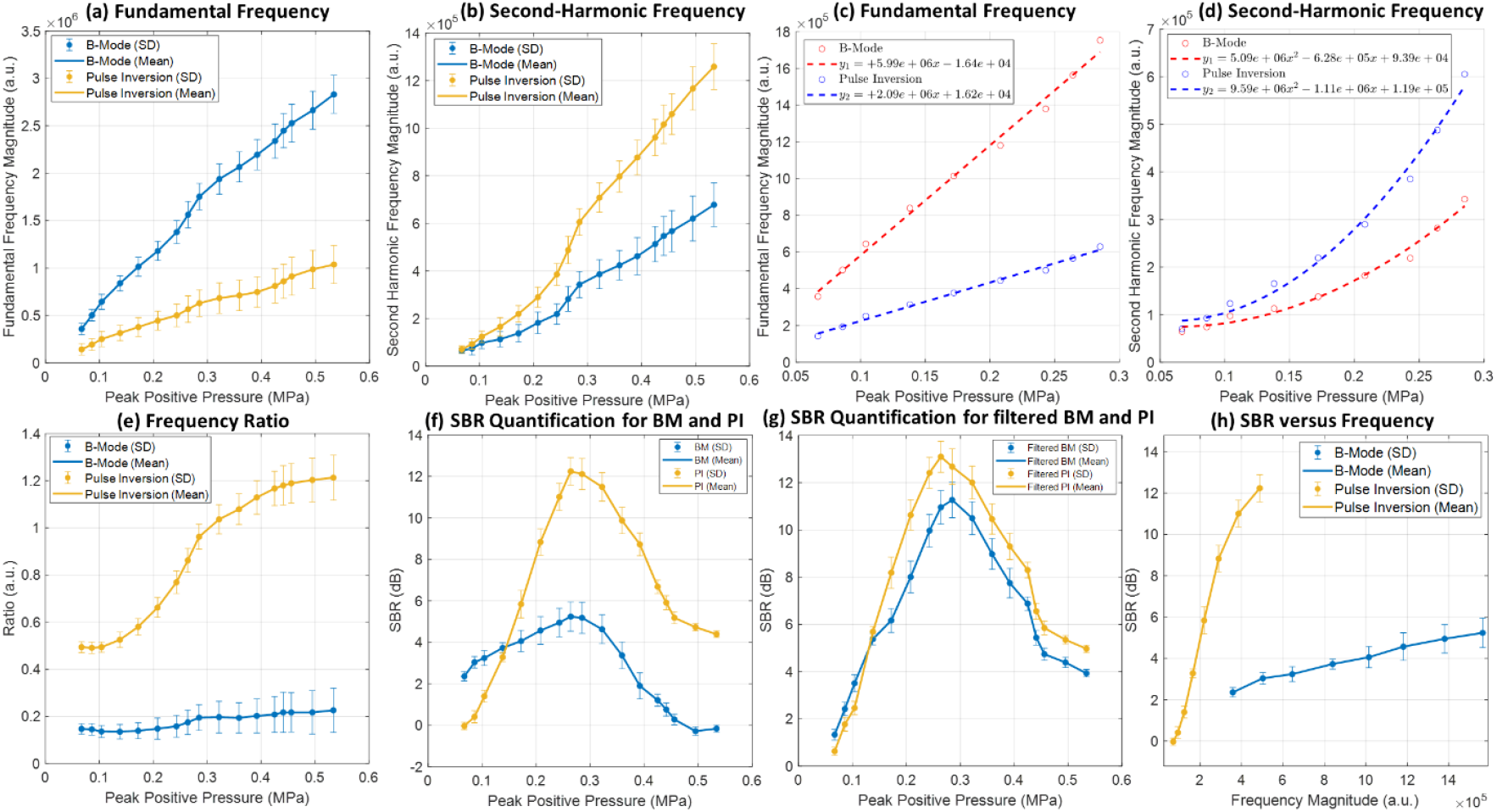
(a) Fundamental and (b) second-harmonic frequency magnitude as a function of the peak positive pressure from 0.067 MPa to 0.534 MPa; (c) Fundamental and (d) second-harmonic frequency magnitude as a function of the peak positive pressure from 0.067 MPa to 0.285 MPa before the GV destruction; (e) the ratio between fundamental and second-harmonic frequency magnitude as a function of peak positive pressure for B-mode and pulse inversion images; SBR quantification for B-mode and pulse inversion images (f) without and (g) with second-harmonic frequency filtering; (h) SBR quantification as a function of fundamental frequency magnitude for B-mode image and second-harmonic frequency magnitude for pulse inversion image.

### 3.3. Simulation

The simulation results presented in Figure 5 present the comparison of simulation between AM-SVD and HAM-SVD on the same numerical gas vesicle phantom, when the number of acoustic pressure level equals to 4 and 16, with the corresponding Fourier transform of pressure singular vectors, and spatial similarity matrix. It can be seen that, when the number of acoustic pressure level equals to 4, both methods showed comparable spatial singular vector distributions, as well as the corresponding Fourier transform of the pressure vectors, spatial similarity matrix, and square fitting matrix. However, when the number of acoustic pressure levels equals to 16, for AM-SVD, GV contrast was mainly distributed from vector 2 to vector 6, while for HAM-SVD, GV contrast was distributed from vector 2 to vector 14, as can be observed from the spatial singular vectors. In terms of spatial similarity matrix, it can be seen that, HAM-SVD has clearer and more uniform block-wised patterns, compared to AM-SVD. This clearer definition of block-wised patterns should provide an easier way to automatically select the singular vectors corresponding to GVs.

**Figure 5.**
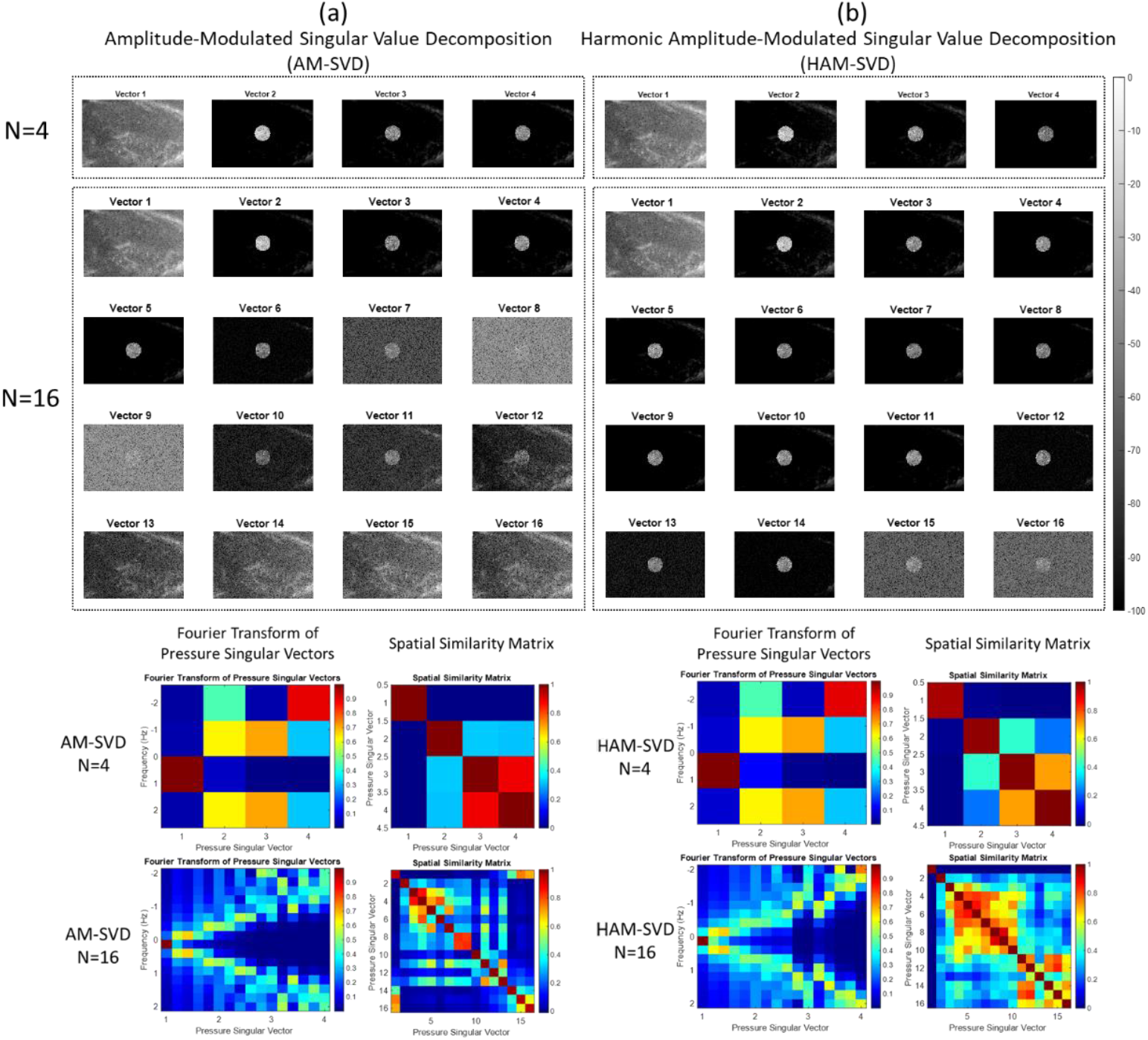
Comparison between AM-SVD and HAM-SVD on the same simulated gas vesicle numerical phantom. (a) Spatial singular vectors of AM-SVD when the number of acoustic pressure level equals to 4 and 16, with the corresponding Fourier transform of pressure singular vectors, and spatial similarity matrix; (b) Spatial singular vectors of HAM-SVD when the number of acoustic pressure level equals to 4 and 16, with the corresponding Fourier transform of pressure singular vectors, and spatial similarity matrix. All the images of spatial singular vectors were displayed in grayscale and in -100 dB.

### 3.4. In Vitro Experiments

Both AM-SVD and HAM-SVD were processed on the same in vitro GV phantom dataset. For the AM-SVD sequence, only the positive frames were used in the processing while for the HAM-SVD sequence, the positive frames and the corresponding negative frames with the identical magnitude were used as input for the SVD processing. As can be seen from Figure 6 (a-b), for HAM-SVD, the first spatial vector demonstrated significantly smaller tissue background signals compared to conventional AM-SVD as the pulse inversion transmission scheme introduced in HAM-SVD already cancels a non-negligible part of the fundamental linear tissue signal. For AM-SVD, some nonlinear image artifacts can be seen below the GV contrast on the spatial vector 2 when N=4, the spatial vectors 2-5 when N=8 and 12. However, the nonlinear image artifacts were not observed in any of the spatial vectors in HAM-SVD. The comparison of SBR quantification between AM-SVD and HAM-SVD in Figure 6(c-e) also shows that the GV contrast was highly concentrated on the first few vectors. Furthermore, HAM-SVD demonstrated higher contrast compared to AM-SVD. This *in vitro* phantom result shows consistency with the simulation results, especially in the spatial similarity matrix of HAM-SVD when N=16. It can be seen that the GV contrast was highly concentrated on the first few spatial vectors, whereas for the AM-SVD, the GV contrast was separated more evenly into different spatial vectors. This indicates that HAM-SVD requires less pressure levels compared to AM-SVD.

**Figure 6.**
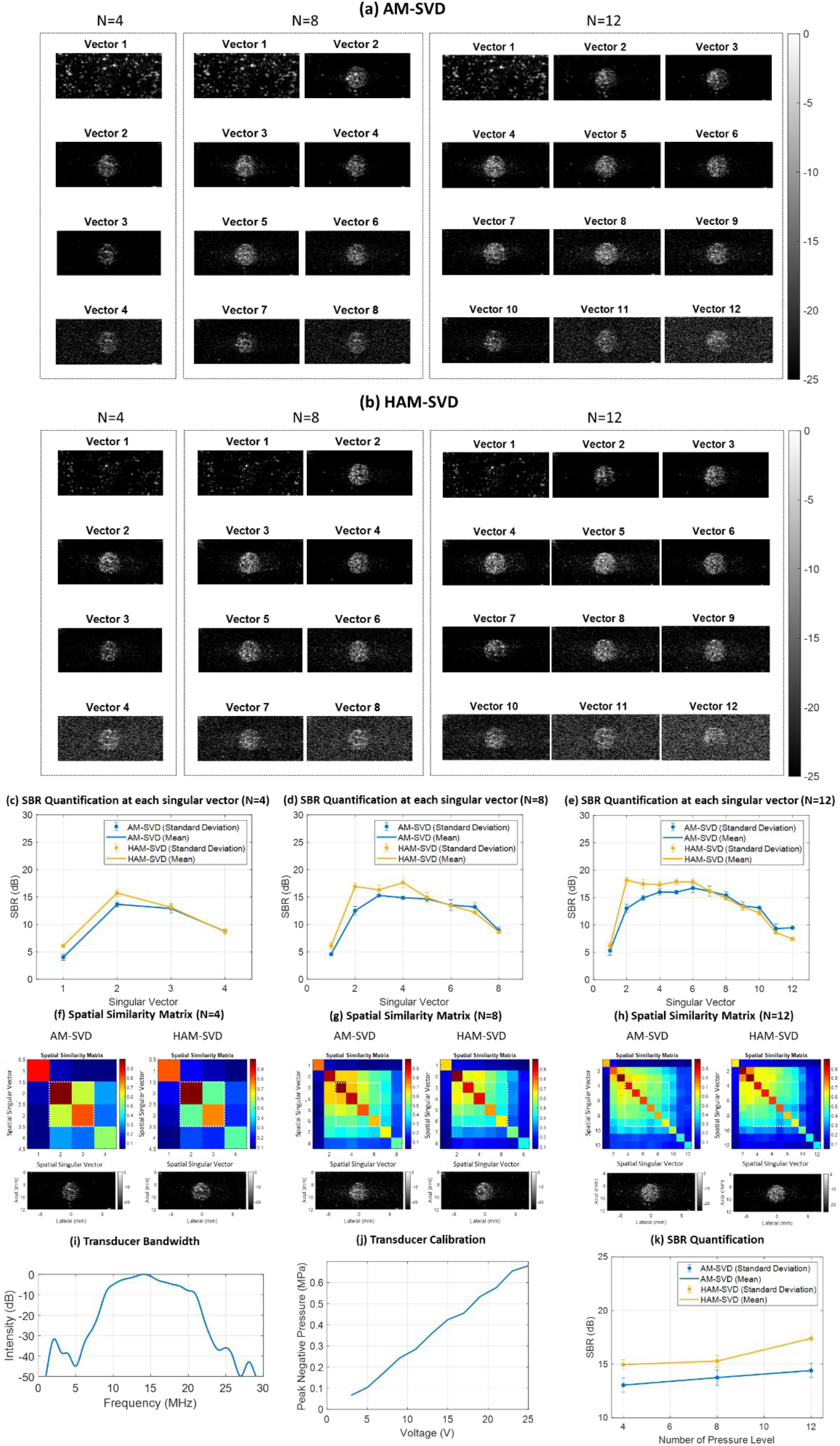
Comparison of *in vitro* experiments between AM-SVD and HAM-SVD on the same gas vesicle phantom. (a) Spatial singular vectors of AM-SVD when the number of acoustic pressure level equals to 4, 8, and 12. The color bar represents the contrast signal in dB; (b) Spatial singular vectors of HAM-SVD when the number of acoustic pressure level equals to 4, 8, and 12. The color bar represents the contrast signal in dB;; (c-e) Quantification of SBR at each singular vector when the number of acoustic pressure level equals to 4, 8, and 12; (f-h) Comparison of spatial similarity matrix between AM-SVD and HAM-SVD. (i) Probe bandwidth (128 elements, 15.625 MHz central frequency, and a 67% bandwidth at -6 dB). The pulse inversion transmission scheme enables to transmit at 9.6 MHz and receive at 19.2 MHz; (j) Probe calibration for the transmission frequency at 9.6 MHz at a depth of 9 mm; (k) Quantification of the final images using different number of pressure levels.

### 3.5. Frequency Spectra of Spatial Singular Vector during HAM-SVD

Figure 7 (a) shows the comparison of 16 spatial singular vectors between AM-SVD and HAM-SVD when the number of pressure levels equals to 16. Figure 7 (c) shows how the frequency spectrum evolved as the spatial singular vector increases using AM-SVD. It can be seen that, for the first three spatial singular vectors, the fundamental frequency components were dominant compared to the second harmonic frequency components. For the later spatial singular vectors, second harmonic frequency components were shown to be higher, which also indicated that, GV contrast was extracted from these spatial singular vectors in AM-SVD. Figure 7 (d) shows how the frequency spectrum evolves as the spatial singular vector increased using HAM-SVD. For the spatial singular vectors from 8 to 11, the ratio between the second harmonic and fundamental frequency peaks were about 2 times.

**Figure 7.**
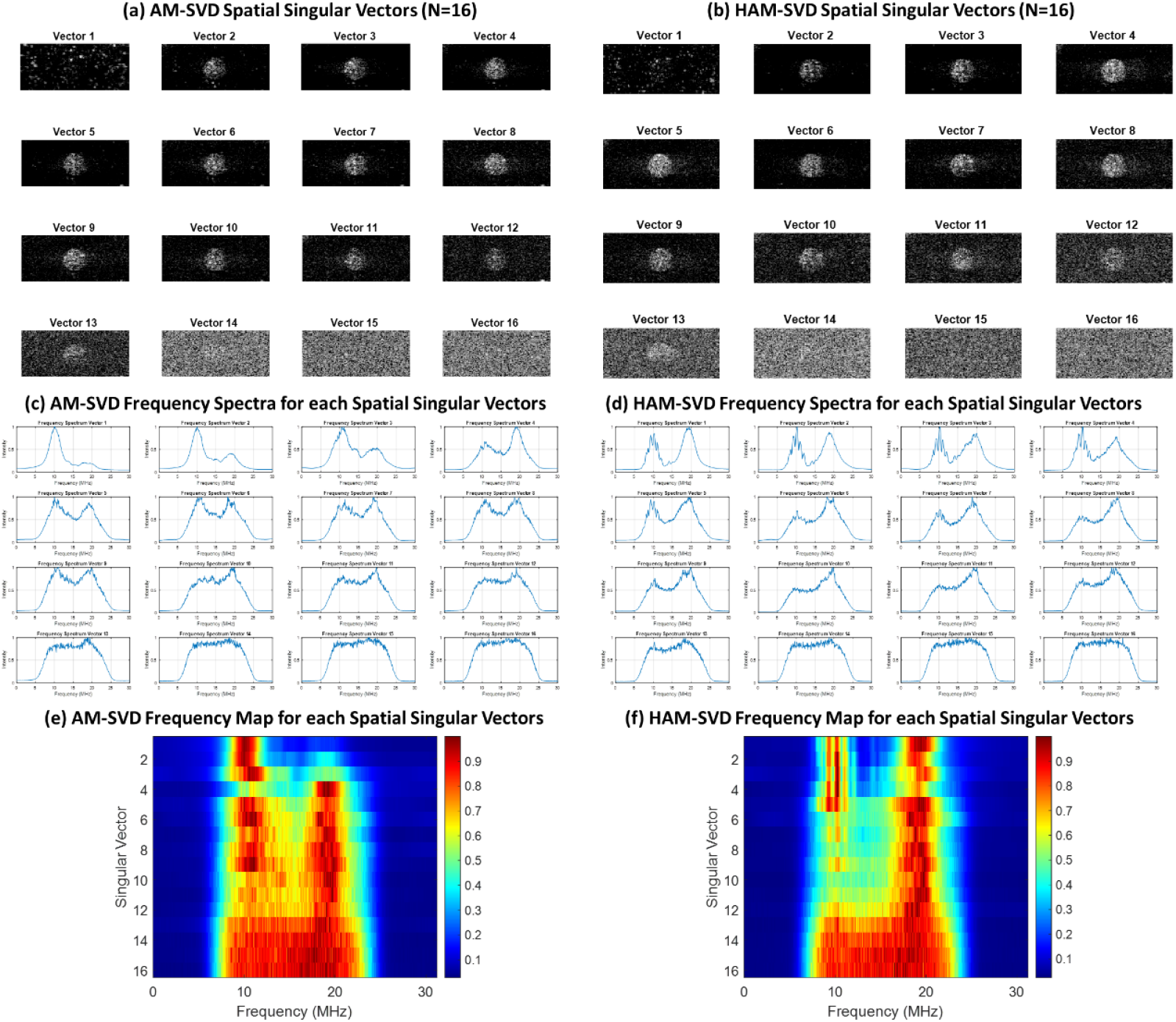
Comparison of spatial singular vectors between AM-SVD and HAM-SVD when the number of pressure levels equals to 16. All of the spatial singular vectors of (a) AM-SVD and (b) HAM-SVD; Frequency spectrum for each spatial singular vectors in (c) AM-SVD and (d) HAM-SVD; Map of frequency distribution as a function of singular vectors using (e) AM-SVD and (f) HAM-SVD.

### 3.6. In Vivo Experiments

Figure 8 presents the *in vivo* comparison between pulse inversion, AM-SVD and HAM-SVD on the same dataset acquired in the rat lower limb. The image comparison between pulse inversion, AM-SVD and HAM-SVD at each peak positive pressure can be found in Supplementary Figure 1. As can be seen from Figure 8 (a) that, the maximum SBR that can be achieved for pulse inversion, AM-SVD, HAM-SVD are 14.19 ± 1.41 dB, 15.79 ± 1.38 dB, and 19.16 ± 1.63 dB, respectively. At relatively low peak positive pressure range (0.138 – 0.359 MPa), conventional pulse inversion provides higher SBR than HAM-SVD. HAM-SVD provides higher SBR at relatively higher pressure range (0.359 – 0.441 MPa) before GV destruction, and for the pressure range (0.441 – 0.768 MPa) after GV started being destroyed.

**Figure 8.**
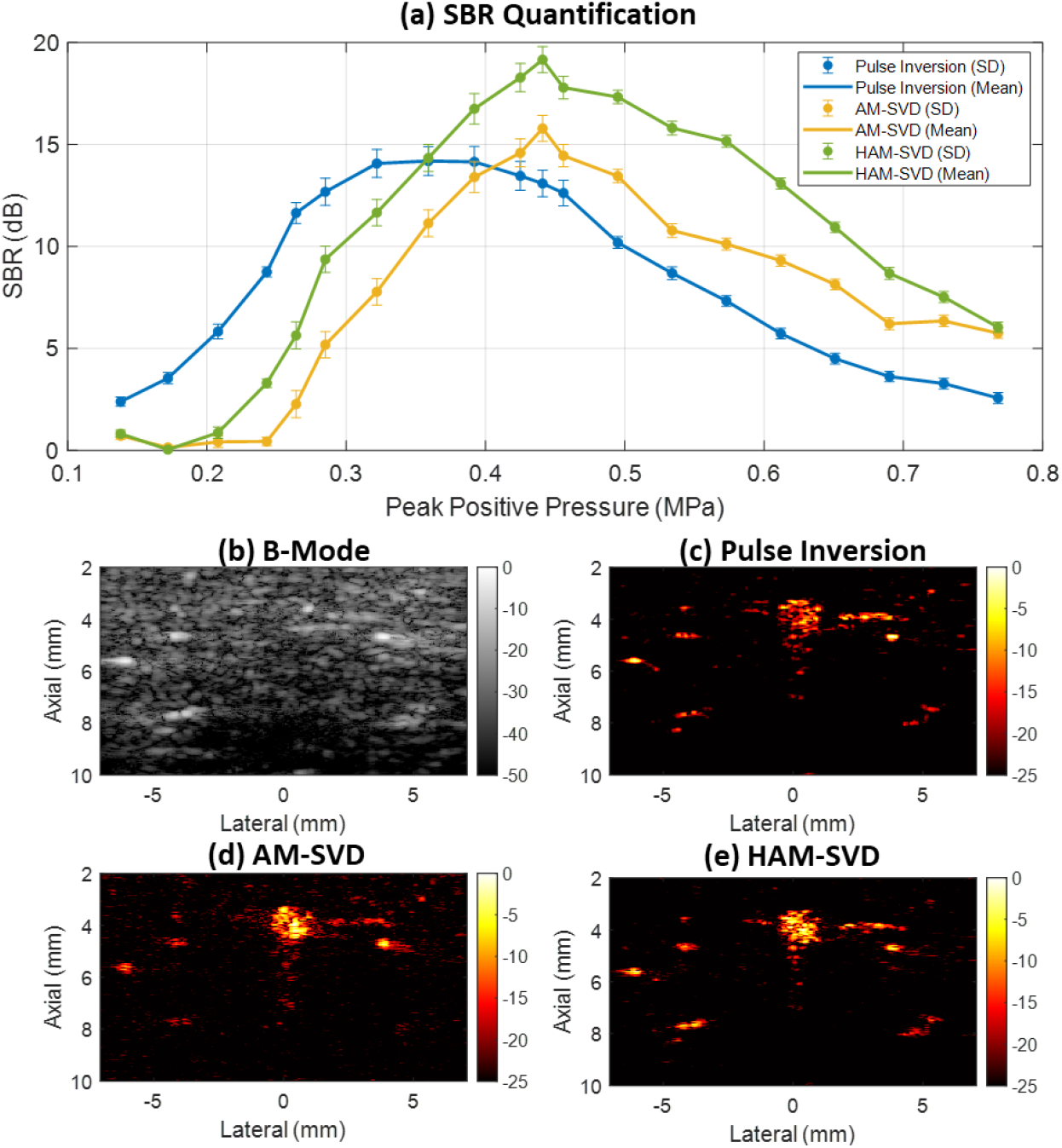
Comparison between conventional B-mode, Pulse Inversion imaging, AM-SVD, HAM-SVD imaging on the same dataset acquired after GV injection in the rat lower limb across the acoustic pressure range of 0.138 – 0.768 MPa. (a) SBR quantification of pulse inversion, AM-SVD, HAM-SVD on the same dataset; (b) B-mode image; (c) Pulse inversion; (d) AM-SVD; (e) HAM-SVD.

## 4. Discussion

This study introduces nonlinear amplitude-modulated singular value decomposition (HAM-SVD) as a novel ultrafast ultrasound imaging technique that exploits the synergetic abilities of pulse inversion (PI) and amplitude-modulated singular value decomposition (AM-SVD) to enhance the detection of gas vesicle contrast agents. By exploiting the distinct second harmonic signatures of GVs compared to tissue signals and leveraging singular value decomposition to isolate pressure-dependent subspaces, this new ultrasound sequence is based on ultrafast plane wave compounded data acquired at different pressure amplitude levels and alternating polarities. The SVD processing applied to these backscattered ultrasound signals comprising the nonlinear signature of GVs and nonlinear propagation in surrounding tissues extracts independent images, each exhibiting a different nonlinear dependence with respect to the transmit amplitude. The cancellation of high-order singular value components permits to filter the signals backscattered by tissues, even when they exhibit a nonlinear pressure dependency, and to select preferentially the nonlinear signals originating from GVs. It is anticipated that our proposed nonlinear SVD beamforming technique will enhance GV contrast detection and suppress tissue background signals, particularly in scenarios involving strong nonlinear tissue signals and the absence of motion of GVs. Furthermore, HAM-SVD is envisioned to be applicable for the enhanced detection of slowly moving microbubbles in Ultrasound Localization Microscopy.

The superior performance of HAM-SVD arises from two aspects – pulse inversion transmission and AM-SVD processing. AM-SVD works on the fundamental frequency, while HAM-SVD works on the second harmonic frequency. It was found in this study that, in the frequency domain of the nonlinear second harmonic, although PI scheme cannot completely remove the tissue signal, it helps remove most of the tissue signals, and thus the remaining tissue signal can be more easily filtered out by HAM-SVD. Additionally, compared to the fundamental signal of GVs, the second harmonic signals of GVs exhibit a stronger nonlinearity with respect to the increase of acoustic pressures compared to tissue signals as reported in the previous literature [6]. For this reason, HAM-SVD sequence can even more easily separate the remaining nonlinear tissue signal and GV signals. These two mechanisms enable HAM-SVD to perform better in the nonlinear second harmonic domain.

It can be seen from Figure 8(a) that PI showed a significantly lower SBR compared to AM-SVD and HAM-SVD when the peak positive pressure is above around 0.35 MPa. There are two possible explanations for this. Firstly, at relatively high pressure, the positive and negative pulses cannot cancel out completely due to an increasing nonlinearity in electronic transmission. Second, at relatively high pressure, the tissue itself will also generate a nonlinear signal on PI imaging. It can also be seen from Figure 8(a) that, PI demonstrated a higher SBR compared to AM-SVD and HAM-SVD, this may be due to the reason that, the differences of nonlinear responses of GVs across 4 different duty cycles were limited at relatively low pressures (eg, below 0.35 MPa).

It can be seen from the *in vivo* results that the GV destruction happened at 0.441 MPa, which differs from 0.6 MPa in the previous literature. The potential explanation could be that, for the most of previous GVs imaging literature, 15 MHz was used as the transmission frequency. However, in our study, the transmission frequency was set as 9.6 MHz in order to be able to receive the second harmonics within the bandwidth of the probe. This suggested that, buckling and collapse pressure of GVs may also be dependent on the ultrasound transmission frequency. In this case, mechanical index (MI) may be a more suitable index to evaluate the GV destruction limit, as MI takes into account the value of transmission frequency. After the MI conversion, the corresponding destruction MIs are 0.14 (for 0.441 MPa at 9.6 MHz) and 0.15 (for 0.6 MPa at 15.625 MHz).

It should be noted that, although AM-SVD only utilized B-mode image on the fundamental frequency domain, it can still extract the second harmonic information during the decomposition, which can be seen from Figure 7 (c). For HAM-SVD, since the input was the pulse inversion images and the fundamental tissue clutter cannot be completely removed in pulse inversion, the fundamental frequency information corresponding to the nonlinear tissue dependency was still retained in the first spatial singular vector, which can be observed in Figure 7 (b).

A limitation of HAM-SVD in this study is that only stationary GV signal was investigated. The performance of HAM-SVD on flowing contrast agents, e.g., microbubble contrast agents in the blood flow, is still unknown. Future studies should also investigate the performance of HAM-SVD on flowing microbubbles. The ability of HAM-SVD to discriminate contrast agents from tissues without exploiting the temporal coherence of signals could provide a first beneficial step before exploiting the clutter SVD filter based on temporal coherence. This two-step filtering, HAM-SVD followed by clutter filter SVD should help to detect more slowly-flowing microbubbles in tiny vessels. Moreover, real-time GPU implementation of HAM-SVD will be essential for a wide range of preclinical translations. Finally, although demonstrated here on purified GVs, HAM-SVD could be extended to *in vivo* imaging of cells expressing GVs for molecular ultrasound imaging.

## 5. Conclusion

HAM-SVD represents a significant advance in ultrafast nonlinear ultrasound imaging by combining pulse inversion and amplitude-modulated singular value decomposition to achieve high-contrast, wide-field visualization of gas vesicles. Through simulations, phantom experiments, and *in vivo* validation, we show that HAM-SVD dramatically enhances GV signal-to-background ratio while reducing artefacts. Its flexibility in pressure selection, its compatibility with ultrafast plane-wave acquisition, and its improved capacity for automated SVD thresholding position HAM-SVD as a powerful tool for preclinical molecular imaging. Ongoing work will focus on real-time implementation and extension to dynamic flow scenarios, paving the way toward preclinical deployment for biomarker detection.

## Acknowledgements

This work was supported by Inserm research accelerator (Inserm ART) in Biomedical Ultrasound and by the French national research agency (ANR) under ANR-21-CE19-0050 program (Project SonoGT). MGS is an investigator of the Howard Hughes Medical Institute. This research was also funded by the Region Ile de France - Convention DIM ELICIT Innovative Technologies for Life Science. This work was also partially funded by the AXA research fund (Project NeuroElastoFlow).

---

**Supplementary Figure 1.**
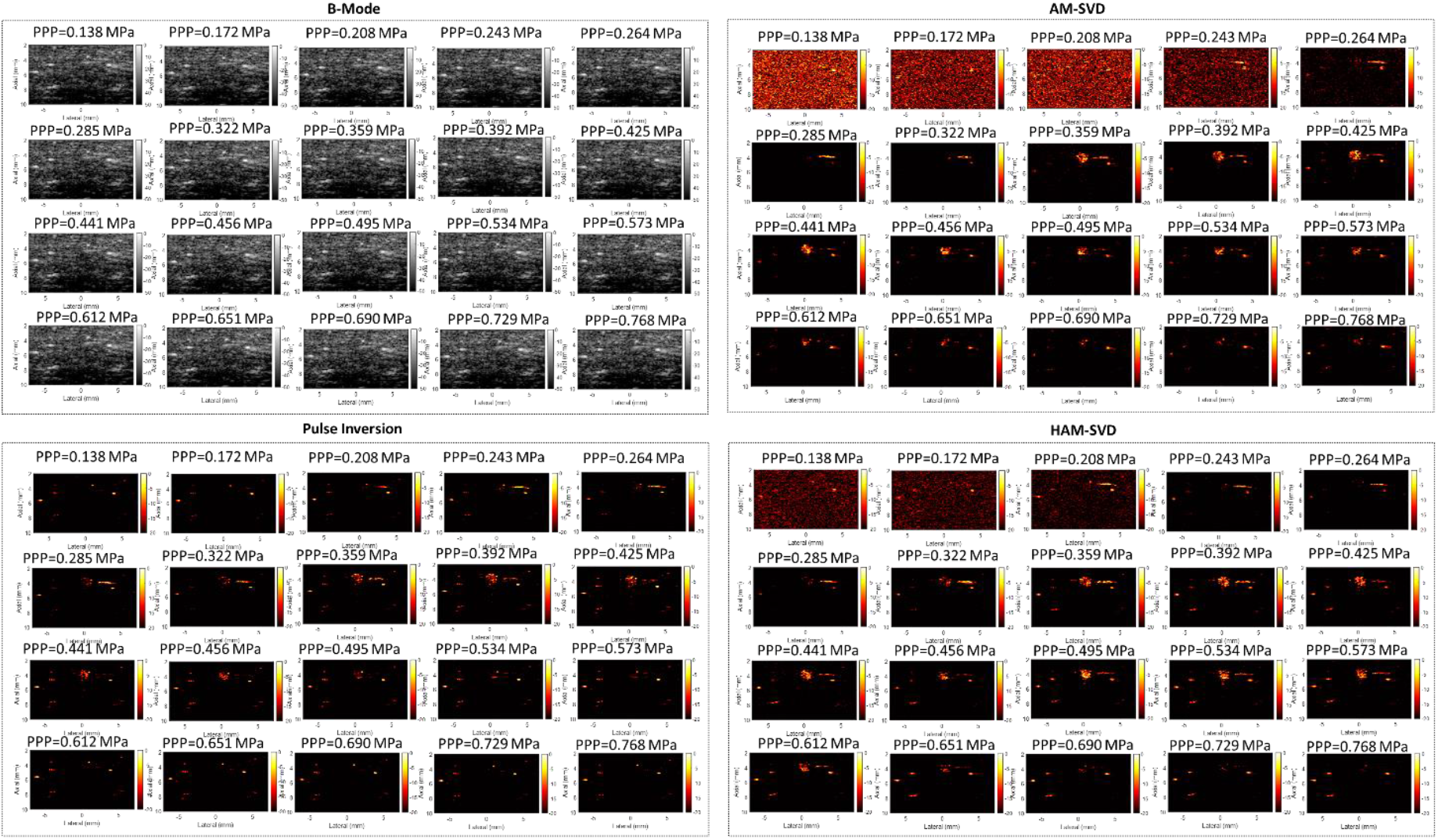
The comparisons between B-mode, pulse inversion, AM-SVD, HAM-SVD on the same dataset acquired on the GV injection on the lower limb of the rat across the acoustic pressure range of 0.138 – 0.768 MPa.

## Supplementary Material 1

### Theoretical frame rate analysis

For a 10mm depth acquisition and number of pressure amplitude N=4 (achieved by 4 different duty cycles), theoretically acquired along the 128 elements of a linear array transducer. One single plane-wave acquisition is then acquired in:

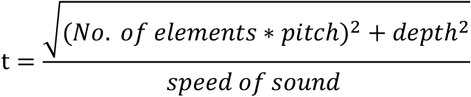

with number of elements = 128, pitch = 0.10 mm, depth = 10mm, and speed of sound = 1540 m/s

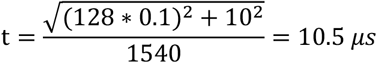

then, one image of the proposed HAM-SVD method (plane-wave transmissions across 11 angles at 4 different pressure amplitudes) and 2 different polarities can theoretically be acquired in:

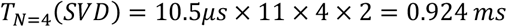

Which corresponds to an imaging frame rate of 1.08 kHz. (As it can be achieved by using a single voltage, therefore, there is no need to change TPC profile in the Verasonics system)

## Notes

### Competing Interest Statement

The authors have declared no competing interest.

